# A Set of Novel Far-Red Fluorescent Proteins for Temporal Domain Multiplexing and Super-Resolution Imaging

**DOI:** 10.1101/2025.06.01.657196

**Authors:** Fakorede Olumayowa, Zhien Rong, Ruizhao Wang, Stavrini Papadaki, Xinyue Wang, Jiayue Cao, Fedor V. Subach, Reinhard W. Köster, Kazuhiko Namikawa, Kiryl D. Piatkevich

## Abstract

The diverse spectral and photochemical properties of fluorescent proteins enable a variety of imaging applications in cell biology, ranging from cellular and organelle labeling to super-resolution microscopy and multiplexed live cell imaging. Here, we report a set of novel far-red fluorescent proteins, named mfRFP, mfRFP-A, and mCardinal-A, which are characterized by similar fluorescence spectra with excitation/emission at ∼600/660 nm while exhibiting distinct photobleaching rates. Differences in photostability allowed us to perform per-pixel unmixing of the three far-red FPs imaged simultaneously by employing a recently introduced temporal domain multiplexing approach. We demonstrated the application of the temporal domain multiplexing approach with different combinations of far-red fluorescent proteins possessing nearly identical emission spectra by acquiring BrainBow-like images of cellular populations and distinguishing subcellular structures in mammalian cells using a single imaging channel without applying any hardware modifications to the conventional microscope. Unlike previous temporal domain multiplying approaches employing photophysical properties of fluorescent proteins, the current approach is a wide range of microscopy modalities, including 3D imaging with a spinning disk and point scanning confocal microscopy. The most photostable fluorescent protein in the set, mfRFP, was further benchmarked against spectrally similar FPs and applied for super-resolution imaging of structural proteins in mammalian cells and for neuroimaging of model organisms, including mice, zebrafish, and *C. elegans*.

## INTRODUCTION

Fluorescence microscopy has driven many prominent breakthroughs in life sciences and continues to play a significant role in new biological discoveries.^1^ Fluorescent proteins (FPs) represent a crucial extension of fluorescence microscopy hardware as they enable real-time, non-invasive visualization of desired cellular structures and proteins through specific labeling on the genetic level.^2,3^ With the rapid advancement of optical hardware, multicolor imaging has become one of the major trends in microscopy, allowing simultaneous observations of several different proteins of interest in individual cells, thus revealing the intrinsic relationships between their subcellular locations and dynamics.^4,5^ Traditionally, signal separation relies on differences in fluorescence spectral profiles or fluorescence lifetimes between FPs.^6^ However, spectral separation typically requires a difference of more than 50 nm between fluorescence spectra. Spectral multiplexing of FPs is fundamentally constrained by the narrow range of the visible spectrum and is technically limited by the finite number of optical filter sets available on standard imaging systems. While it is possible to unmix spectrally similar FPs, this process demands the precise design of optical filters and meticulous selection of suitable FPs based on their exact fluorescence spectra.^7,8^ These complexities have hindered the widespread adoption of this approach by average users. In contrast, fluorescence-lifetime imaging microscopy (FLIM) is a well-established technique for multiplexed imaging of spectrally similar fluorescent proteins (FPs)^9^ However, due to the complexity and expense of the FLIM hardware, it remains relatively inaccessible. Furthermore, achieving effective multicolor imaging with FLIM requires substantial differences in fluorescence lifetimes and adequate brightness of used FPs.

More recently, two alternative approaches called the extended intra-exposure excitation modulation method (exNEEMO)^10^ and temporally multiplexed imaging (TMI)^11^ were successfully employed for signal separation of up to 6 FPs in mammalian cells using just one spectral channel. Both methods rely on the reversible photoswitching of GFP-like FPs characterized by distinct photoswitching kinetics to generate brief time-lapse imaging datasets. Acquired datasets are then processed to calculate a weighted sum of the reference traces from each expressed fluorophore in each pixel. Reference traces are recorded in a separate experiment when only one FP is expressed and imaged under matching conditions. The weighted contributions of each FP in a pixel are presented as intensities in different channels corresponding to FPs reconstructing a quantitative multicolor image. As a result, a series of images acquired in a single spectral channel over a time period required to photoswitch FPs is converted into a multicolor image, where each color represents a defined FP. The ExNEEMO and TMI methods, which we refer to as temporal domain multiplexing (TDM) in this study for simplicity, represent several advantages over existing traditional methods for multiplexed images, such as multispectral imaging and fluorescence lifetime imaging microscopy (FLIM). For example, in contrast to FLIM, which requires the addition of expensive and sophisticated equipment to an imaging system, TDM can be performed on conventional microscopes commonly available to biologists. Furthermore, TDM can extend the multiplexing capabilities of the existing imaging setups by enabling imaging of multiple FPs in one channel. However, to date, only green and red reversible photoswitchable FPs were applied for TDM. Moreover, due to fast photoswitching, on the order of a few seconds, used FPs are not feasible for 3D multiplexed imaging or certain imaging modalities, such as point scanning confocal microscopy. Altogether, these issues further limit the applicability of TDM for cell biology imaging.

We sought to extend the applicability of TDM beyond reversibly photoswitchable FPs and test whether multiplexing can be done with conventional or so-called permanently emitting FPs. In this case, signal separation can be achieved through differential photobleaching rates of spectrally similar FPs. Since green and red spectral ranges have already been used for TDM with exNEEMO and TMI, we chose to use far-red FPs, which possess an additional advantage over green and red FPs due to the lower phototoxicity of the corresponding excitation light. The latter is particularly important as photobleaching of FPs may require extended illumination that may cause undesired phototoxicity to cells. In this study, we report a set of novel far-red fluorescent proteins, namely mfRFP, mfRFP-A, mCardinal-A, characterized by distinct photobleaching rates in the range of excitation powers from 10 to 100 mW/mm^2^ typically used for live cell imaging. The mfRFP, mfRFP-A, and mCardinal-A proteins were carefully characterized in solution and benchmarked against spectrally similar FPs in live and fixed mammalian cells. Using different combinations of the developed FPs, we were able to successfully perform 3D TDM imaging of different structural proteins in single cells as well as BrainBow-like imaging of cellular populations in culture. Due to the superior photostability of mfRFP, we further demonstrated its applicability for STED microscopy and neuroimaging in mice, zebrafish, and nematodes. We also discovered the reversible photoswitching behavior of mfRFP-A with on/off switching induced by 488/594 nm, which makes it the first photoswitchable far-red FP. We anticipate that the newly introduced set of far-red FPs will be a valuable addition to the cell biology toolset, particularly for multiplexed imaging and advanced microscopy.

## Results

### Development and characterization of the novel far-red FPs

To develop far-red FPs with distinct photobleaching rates, we employed a directed molecular evolution approach^12^ to diversify the photobleaching rates of the recently introduced TagRFP658 protein characterized by excitation/emission at ∼611/658 nm^13^. Bacterial libraries generated using error-prone PCR were subjected to screening with a fluorescence-activated cell sorter (FACS), followed by a selection of the brightest clones on Petri dishes using a fluorescence stereomicroscope. For screening, we used 637 nm excitation and a 660 nm long-pass filter for fluorescence emission to maintain selection pressure on red-shifted mutants. Using shorter excitation wavelengths could bias the selection toward blue-shifted variants, as these mutants are typically brighter. The 10–15 brightest clones in each round were assessed for photostability under continuous wide-field excitation, and the most and least photostable variants were subjected to the next round of evolution. As a result, after several rounds of evolution, we selected two variants, named mfRFP and mfRFP-A, exhibiting more than a 10-fold difference in photostability under screening conditions. The mfRFP and mfRFP-A proteins differed by a single amino acid substitution (T163A) from each other while by 8 and 9 mutations, respectively, from the parental TagRFP658 protein (**Supplementary Figure 1**). Suggesting that T163A was responsible for reduced photostability, we utilized mCardinal, one of the brightest far-red FPs, as a template to introduce identical amino acid at the corresponding position with the purpose of generating its less photostable variant, which we refer to as mCardinal-A (**Supplementary Figure 1**; see the results of photostability assessment below).

We proceeded with spectral and biochemical characterization of the generated variants in the solution. For comparison, we also purified and characterized one intermediate variant of mfRFP, mfRFP0.1, and four other non-obligate dimer GFP-like FPs with emission maxima beyond 650 nm, namely mCarmine^14^ mMaroon^15^, mGarnet2 (ref.^16^), mCardinal^17^, which are characterized by the highest molecular brightness among far-red FPs. The mfRFP, mfRFP-A, and mCardinal-A proteins exhibited very similar spectral profiles with main absorbance/excitation maxima in the range of 598-608 nm and emission peaks at around 658 nm (**Figure 1a-c**, **Table 1**). The molecular brightness of mCardinal-A was 1.2-, 2.2-, and 3.2-fold higher than mCardinal, mfRFP, and mfRFP-A, respectively, also surpassing the molecular brightness of other tested far-red FPs, thus making mCardinal-A one of the brightest far-red GFP-like FPs with emission maximum beyond 650 nm (**Table 1**). All variants demonstrated moderate pH stability of fluorescence with pKa values in the range of 4.8 to 5.5, with mfRFP being the most pH-stable FP among all tested FPs (**Figure 1d**, **Table 1**). The emission peak remained unchanged for mfRFP and mfRFP-A; however, for mCardinal and mCardinal-A, it exhibited a 4 nm blue shift at pH levels below 6.0 (**Supplementary Figure 2**). As a result, the generated far-red FPs exhibited higher or comparable molecular brightness and pH stability relative to the state-of-the-art far-red FPs with matching fluorescence spectra.

**Figure 1.**
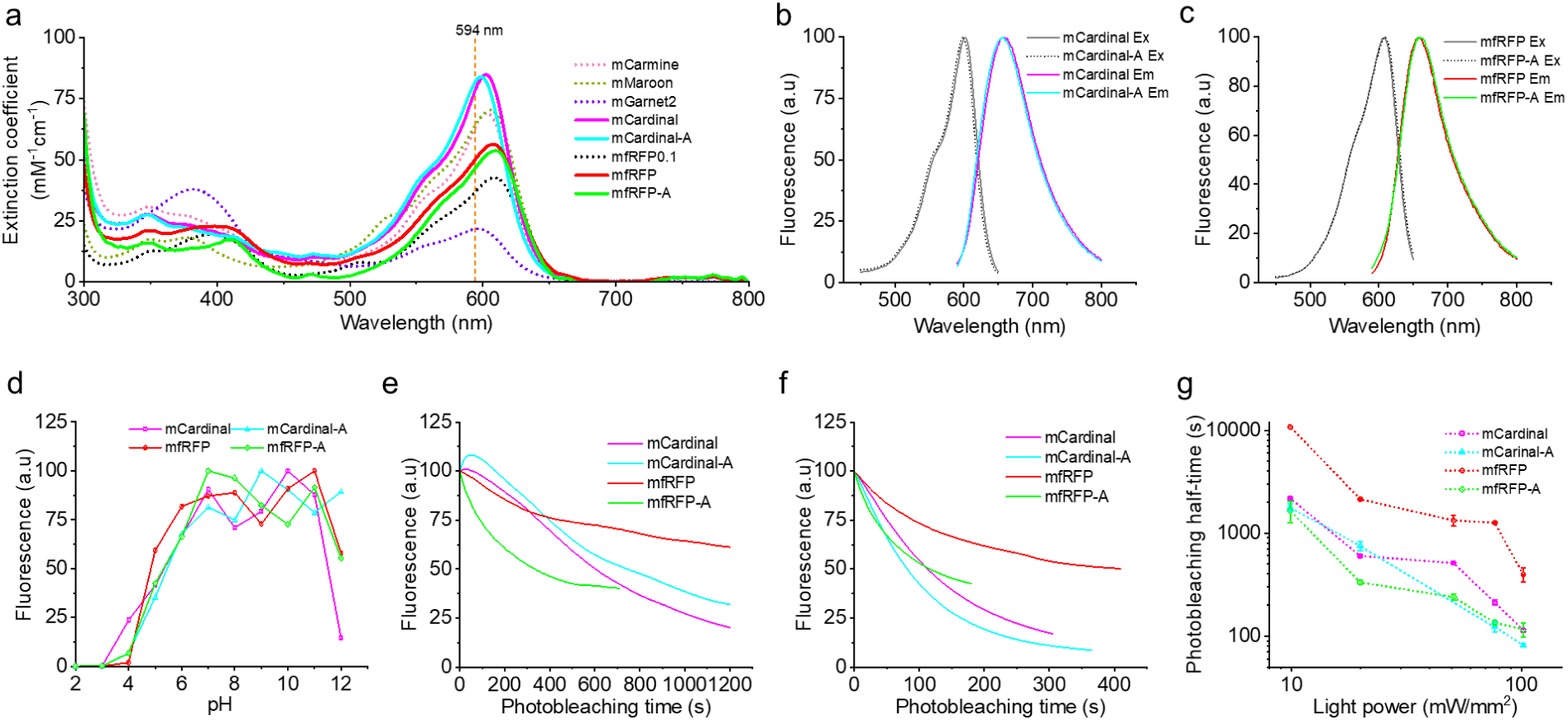
In vitro characterization of the selected far-red FPs in solution. (a) Absorbance spectra of mCarmine, mMaroon, mGarnet2, mCardinal, mCardinal-A, mfRFP0.1, mfRFP, and mfRFP-A. (b) Fluorescence excitation and emission spectra of mCardinal and mCardinal-A. (c) Fluorescence excitation and emission spectra of mfRFP and mfRFP-A. (d) pH dependence of fluorescence for mCardinal, mCardinal-A, mfRFP, and mfRFP-A (one technical replicate each). (e) Photobleaching curves of mCardinal, mCardinal-A, mfRFP, and mfRFP-A droplets in immersion oil under wide-field illumination at 575/50 mn (n = 6, 10, 10, and 10 ROIs, respectively, excitation power 20 mW/mm^2^). (f) Photobleaching curves of mCardinal, mCardinal-A, mfRFP, and mfRFP-A droplets in immersion oil under wide-field illumination at 575/50 mn (n = 10, 10, 10, and 10 ROIs, respectively; excitation power 102 mW/mm^2^). (g) Photostability half-time versus illumination power measured under wide-field illumination at 575/50 nm for mCardinal, mCardinal-A, mfRFP, and mfRFP-A droplets in immersion oils at increasing illumination power (n = 9, 6, 4, 10, 10 ROIs for mCardinal, 10, 10, 0, 10, ROIs for mCardinal-A, 10,10,10,10 ROIs for mfRFP and 10, 10, 10, 10, 10 ROIs for mfRFP-A, at 10, 20, 51, 77,102 mW/mm^2^, respectively). Dot, mean; error bars, SD.

**Table 1.** Spectral and biochemical characterization of far-red FPs in solution and mammalian cells.

| **Protein** | **Abs.**<br>**(nm)** | **Em.**<br>**(nm)** | **EC (M<sup>-1</sup>cm<sup>-1</sup>)** | **QY (%)** | **Molecular**<br>**brightness** | **pK<sub>a</sub>** | **Photobleaching**<br>**half-time (s)** | **Ref.** |
| --- | --- | --- | --- | --- | --- | --- | --- | --- |
| **mCarmine** | 603 | 659 | 70,431±12,820 | 0.1338 | 9423 | 5.6 | 1201 | <sup>14</sup> |
| **mMaroon** | 609 | 657 | 70,859±4,800 | 0.1063 | 4393 | 6.2 | NA | <sup>15</sup> |
| **mGarnet2** | 596 | 674 | 22,070 | 0.056 | 1235 | 6.8 | 1034 | <sup>16</sup> |
| **mCardinal** | 603 | 658 | 85,038±4,638 | 0.1201 | 10213 | 5.3 | 140 | <sup>17</sup> |
| **mCardinal-A** | 598 | 657 | 88,046±2,829 | 0.14 | 12326 | 5.5 | 104 | This study |
| **emiRFP670** | 642 | 670 | 87,400 | 0.14 | 12240 | 4.5 | NA | <sup>36</sup> |
| **TagRFP658** | 611 | 658 | 41,200 | 0.11 | 4532 | 4.6 | NA | <sup>28</sup> |
| **mfRFP0.1** | 609 | 660 | 42,850±2,867 | 0.1394 | 5973 | NA | NA | This study |
| **mfRFP** | 606 | 658 | 57,445±10,079 | 0.0972 | 5583 | 4.8 | 1199 | This study |
| **mfRFP-A** | 608 | 658 | 53,321±10,891 | 0.0719 | 3833 | 5.3 | 41 | This study |

Next, the photostability of the developed proteins was characterized in solution under continuous wide-field illumination. For excitation (here and throughout the study unless otherwise stated), we used a laser or LED with a center wavelength of 594 nm, which efficiently matched the absorption profile of the FPs (**Figure 1a**). As photobleaching rates depend on excitation power^18,19^, we measured photostability at five different light intensities ranging from ∼10 mW/mm^2^ to ∼100 mW/mm^2^, covering typical excitation powers used for cell culture and small model organism imaging. Since the TDM algorithm relies on actual photobleaching curves, we did not perform photobleaching rate correction for molecular brightness and excitation spectrum, as often done for comparison of FPs^19,20^. As expected, photobleaching rates accelerated with increasing illumination power, exhibiting complex supralinear dependence (**Figure 1e-g**). Moreover, at light powers below ∼20 mW/mm^2^, mCardinal, and mCardinal-A demonstrated minor (<10%) initial photoactivation (**Figure 1e**). We found that the high illumination power, *i.e.*, 75-100 mW/mm^2^, would be more suitable for the TDM unmixing as it enabled the acquisition of distinct photobleaching curves in the shorter time scales, *e.g.*, less than 5 minutes at 100 mW/mm^2^ (**Supplementary Figure 3**). Therefore, for all further experiments, we used light powers at around 75-100 mW/mm^2^.

Since our intention was to utilize developed FPs for TDM in mammalian cells, we carried out a pairwise comparison of mCardinal vs. mCardinal-A and mfRFP vs. mfRFP-A as the FPs in these pairs demonstrated matching molecular brightness while distinct photobleaching curves, thus representing potential partners for TDM. Upon transient expression in HeLa cells, all FPs produced easily detectable far-red fluorescence evenly distributed throughout the cells and well co-localized with co-expressed GFP (**Supplementary Figure 4**). The normalized intracellular brightness of mCardinal was 1.37-fold higher than that of mCardinal-A, while mfRFP was only 14% brighter than mfRFP-A (**Supplementary Figure 4**). Photobleaching half-times for mCardinal and mCardinal-A were similar (138 s and 105 s, respectively), though photobleaching curves had distinct shapes (**Supplementary Figure 4**). In turn, mfRFP was more than 8-fold more photostable than mfRFP-A producing drastically different photobleaching curves (**Supplementary Figure 4**). This assessment demonstrated that the developed far-red FPs possessed characteristics suitable for TDM imaging in mammalian cells.

In summary, using directed molecular evolution, we developed a set of novel far-red FPs characterized by distinct photobleaching curves with a more than 10-fold difference in photostability while maintaining comparable spectral properties, molecular brightness, and pH stability to state-of-the-art far-red FPs.

### Temporal domain multiplexed imaging

Having characterized the developed far-red FPs, we proceeded with employing them for TDM imaging of various structures in mammalian cells. TDM imaging involves five major steps: (I) image acquisition; (II) image pre-processing; (III) image unmixing; (IV) output image validation; and (V) recording reference photobleaching curves, as schematically illustrated in **Figure 2a**. Specifically, cultured mammalian cells co-expressing FP fusions were imaged under continuous illumination to reduce initial fluorescence by 50-70%. It typically requires 1-5 min of imaging, depending on the used FPs, illumination power, and sample preparation (fixed or live cells). In the independent experiments, reference traces for the target FPs were recorded under the matching imaging conditions with just one fluorophore expressed at a time (see details in the Methods section). Next, the raw datasets, pre-processed by background subtraction and denoising (optional, used to improve overall image quality), were computationally unmixed using the MATLAB code with input reference photobleaching tracers and then validated with the post hoc image analysis. To evaluate the proposed TDM workflow with far-red FPs, we co-expressed the COX8A-mCardinal and H2B-mfRFP fusions in HeLa cells and recorded time-lapse series to achieve observable photobleaching of at least one FP. Before photobleaching, the selected cell was imaged with a commercially available FLIM scanning confocal system to acquire images for ground truthing the TDM unmixing results. The phasor FLIM plot and reference photobleaching curves exhibited good separation between the mCardinal and mfRFP values, ensuring robust unmixing (**Figure 2b,c**). The time-lapse dataset consisting of 20 frames was processed with the TDM algorithm as described above, and output images were compared to ground truth results obtained using FLIM unmixing (**Figure 2d**). The mCardinal and mfRFP images from TDM showed patterns identical to those generated by the FLIM unmixing, recapitulating the expected cellular morphologies of mitochondria and nucleus, respectively (**Figure 2d**). To quantitatively benchmark TDM unmixing, we calculated the Spearman correlation coefficients between FLIM and TDM images. The Spearman coefficients for line cut intensity profiles from the mCardinal and mfRFP channels were in the range of 0.82-0.89, which was somewhat lower than that between raw FLIM and TDM images (0.96; **Figure 2e,f,g**). We further evaluated FP-FP crosstalk by calculating the fluorescence brightness contribution of FP1 to the FP2 channel and vice versa, following the previously reported approach^21^ (see Methods for details). In FLIM images, mCardinal fluorescence contributed minimally to the mfRFP channel, whereas mfRFP exhibited some cross-talk into the mCardinal channel (**Figure 2h**). In contrast, TDM images showed the opposite trend, with minimal mfRFP cross-talk into the mCardinal channel but a more pronounced contribution of mCardinal fluorescence to the mfRFP channel (**Figure 2i**). Overall, these results demonstrate that TDM can effectively differentiate fluorophores based on their photobleaching characteristics while preserving accurate spatial resolution across different cellular compartments comparable to FLIM. Having validated TDM with FLIM under a point-scanning confocal microscope, we proceeded with testing other imaging modalities, including wide-field and spinning disk confocal microscopy, using various combinations of FPs. For example, wide-field microscopy enabled the separation of mfRFP and mfRFP-A with minimal cross-talk when expressed as H2B fusion in the nucleus and in the cytoplasm, respectively (**Figure 2j,k,l**). Similarly, we successfully unmixed mCardinal and mfRFP-A, expressed in mitochondria and cytoplasm, respectively (**Figure 3m,n,o**). Under spinning disk confocal microscopy, it was possible to unmix a combination of three FPs, namely mRuby2, mCardinal, and mfRFP, exclusively expressed as H2B fusions in HeLa cells (**Figure 2p,q**), although the noticeable contribution of mRuby2 and mfRFP fluorescence into mCardinal channel was observed (**Figure 2r**). This cross-bleed might be due to the lower fluorescence intensity of mCardinal compared to mRuby2 and mfRFP in the raw image (see the raw image in **Figure 2q**).

**Figure 2.**
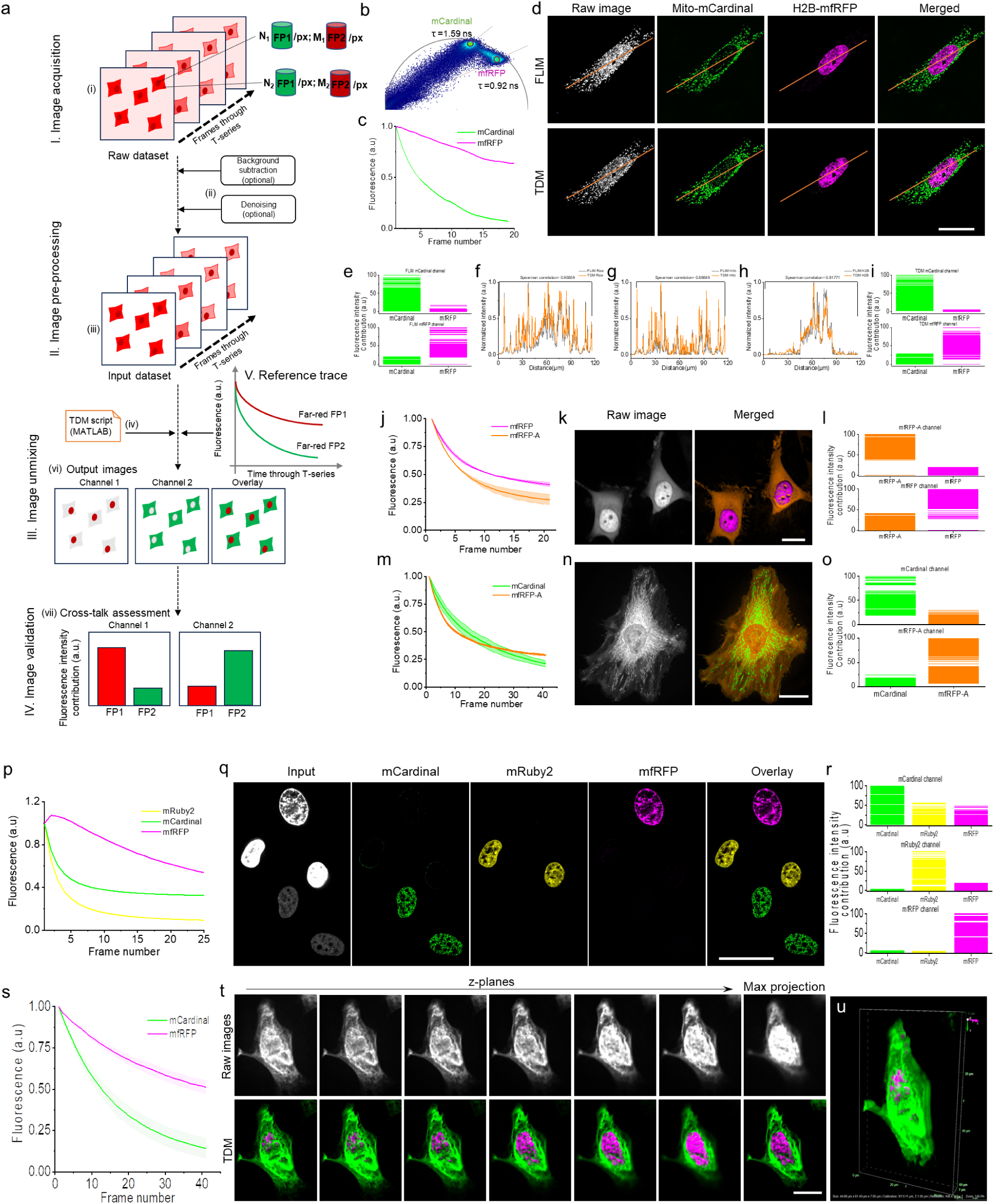
TDM imaging of mammalian cells. (a) Workflow of temporal domain multiplexing, (i) Multi-frame time series acquisition of Hella cell co-transfect with two FPs expressed in the nucleus and cytoplasm, the Raw data set is processed by background subtraction and denoising to produce the input dataset, the input dataset is uploaded to the Temporal domain multiplexing MATLAB script for unmixing to produced output images of individual channels. (b) Phasor plot and lifetime for Mito-mCardinal (green) (1.593ns) and H2B-mfRFP (red) (0.922ns) (c) Reference photobleaching curve trace for Mito-mCardinal (green) and H2B-mfRFP (red). (d) Unmixing results of FLIM microscopy (top), The raw image was unmix using phasor plot analysis resulting in two different FPs Mito-mCardinal (green) and H2B-mfRFP (red) TDM (bottom) unmixing result from the raw image (gray) was supplied to MATLAB script for linear unmixing of each reference at each pixel (the input images are brief movies obtained from single channel imaging containing two different but spectrally similar (far-red) fluorescent proteins. the output image is a composite of Mito-mCardinal (green) and H2B-mfRFP (red). Scale bar 50µm. (e) Cross-talk plot showing the fluorescence intensity contribution of mito (magenta) and H2B (red) in FLIM mito-mCardinal channel (above) and FLIM mfRFP channel (below), in each plot, the y-axis represents the fluorescence intensity of selected pixel, and the x-axis represent the fluorophore in each channel. (f) Per pixel Line profile analysis of FLIM raw image (grey) and TDM raw image (orange) Statistics Spearman correlation (0.95859) the y-axis represents the normalized fluorescence intensity, and the x-axis represent the length (g) Per pixel Line profile analysis of FLIM mito-mCardinal image (grey) and TDM mito-mCardinal image (orange) Statistical Spearman correlation (0.88849) the y-axis represents the normalized fluorescence intensity, and the x-axis represent the length. (h) Per pixel Line profile analysis of FLIM H2B-mCardinal image (grey) and TDM H2B-mCardinal image (orange) Statistics Spearman correlation (0.881771) the y-axis represents the normalized fluorescence intensity, and the x-axis represent the length. (i) Cross-talk plot showing the fluorescence intensity contribution of mito (magenta) and H2B (red) in TDM mito-mCardinal channel (above) and TDM mfRFP channel (below), in each plot, the y-axis represents the fluorescence intensity of selected pixel, and the x-axis represent the fluorophore in each channel. (j) Simulation of Temporal domain multiplexing with two far-red fluorescent proteins, A 21-frame line plot obtained from previously acquired HeLa cell ROI illustrating the unique photobleaching curve of mfRFP and mfRFP-A that contribute to the overall signals, Data are shown as ± mean standard deviation SD. Illumination(594nm at 100% laser power, 500ms Exposure time. The line plot is subsequently used as reference trace for the unmixing algorithms (k)Acquired brief movie (Time series image) of HeLa cell expressing H2B-mfRFP and mfRFP-A-cytoplasm acquired using 594nmSoRA excitation laser channel. The unmixing result (MATLAB) the signal contribution of each FPS (OUTPUT) to their respective corresponding structures and the merged image depicting the complete HeLa cells structure (H2B and cytoplasm). Note that the image colors used above red-H2B and green-cytoplasm are pseudo colors to enhance visualization and both H2B and the cytoplasm were indeed red. Scale bar 25μm. (l) crosstalk plot showing the fluorescence intensity contributions of mfRFP and mfRFP-A in the TDM output channels, the y-axis represents the fluorescence intensity of selected pixel, and the x-axis represent the fluorophore in each channel. (m) Acquired photobleaching reference traces of mCardinal and mfRFP-A in life HeLa-cell, n=5 cells from 2 briefed movies, 41 frames long, from 1 cell culture batch. Data are shown as ± mean standard deviation SD. Illumination(594nm at 100% laser power, 500ms Exposure time (n), The input image was first refined via background subtraction (ImageJ) followed by denoising (Nikon Element AR software), the input image was supplied to MATLAB script for linear unmixing of each reference at each pixel (the input images are brief movies obtained from single channel imaging(red), containing two different but spectrally similar (far-red) fluorescent proteins. the output image is a composite of mfRFP-H2B and Mito-mCardinal. (Scale-Bar 100um). (o) crosstalk plot showing the fluorescence intensity contributions of mCardinal and mfRFP-A in the TDM output channels, the y-axis represents the fluorescence intensity of selected pixel, and the x-axis represent the fluorophore in each channel. (p) Simulation of TDM using Confocal microscope acquired photobleaching reference traces of H2B-mCardinal, H2B-mRuby2 and H2B-mfRFP in fixed HeLa-cell, 25 frames long (q), the input image was supplied to MATLAB script for linear unmixing of each reference at each pixel (the input images are brief movies obtained from single channel imaging(red), containing three different but spectrally similar (far-red) fluorescent proteins of H2B-mCardina, H2B-mruby2 and H2B-mfRFP, and overlay image is a composite. (Scale-Bar 100um). (r) crosstalk plot showing the fluorescence intensity contributions of H2B-mCardinal, H2B-mRuby2 and H2B-mfRFPin the TDM output channels, the y-axis represents the fluorescence intensity of selected pixel, and the x-axis represent the fluorophore in each channel. (s) Temporal domain multiplexing of six z-plain projection of two red fluorescent proteins(FPs). (A) Acquired photobleaching reference traces of mfRFP and mCardinal in life HeLa-cell, n=10 cells from 2 briefed movies, 41 frames long, from 1 cell culture batch. Data are shown as ± mean standard deviation SD. Illumination(594nm at 100% laser power, 500ms Exposure time). (t) Demultiplexing of co-expressed Keratin-targeted mCardinal and H2B-targeted mfRFP in HeLa cell. the raw image (top) was supplied to MATLAB script for linear unmixing of each reference at each pixel, the Merge TDM image (bottom) is a composite of mfRFP-H2B and mCardinal-Keratin. The output image mfRFP-H2B and mCardinal-Keratin are products of linear un-mixing (Scale-Bar 100um) (u) 3D image obtained from the z-stack of the unmixed z-planes images.

**Figure 3.**
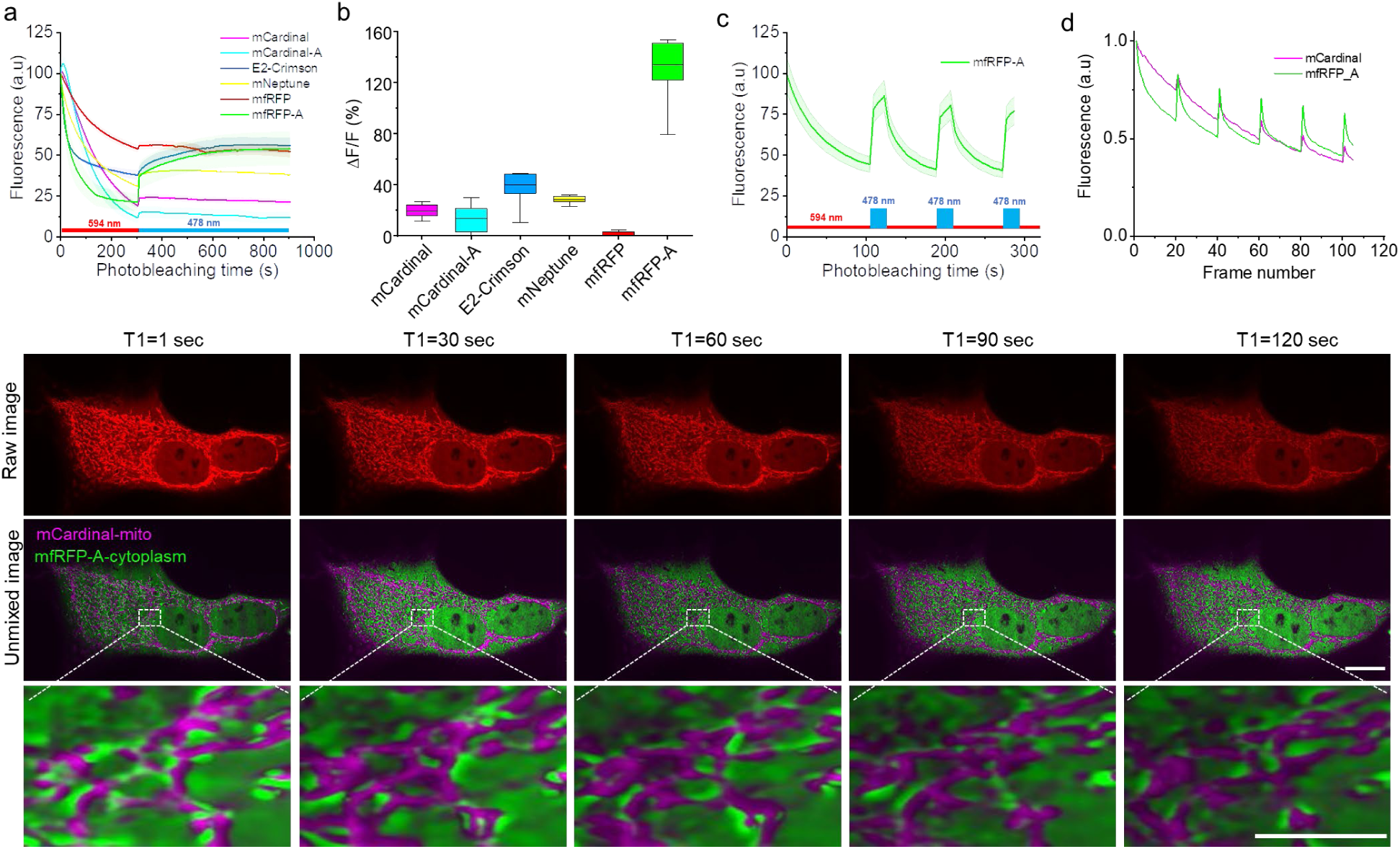
Reversible photoswitching of mfRFP-A enables TDM in live cells to track mitochondria dynamics. (a). Far-red fluorescence traces of mCardinal mCardinal-A, E2-Crimson, mNeptune, mfRFP, and mfRFP-A, under continuous 594 nm excitation followed by induction of photoreversibility using the 488 nm excitation for 10 minutes. (b). Bar plot with SD (whisker) depicting the amplitude of change of intensity of selected far-red fluorescent proteins mCardinal, mCardinal-A, E2-Crimson, mNeptune, mfRFP and mfRFP-A with induction of photo-reversibility with 488 nm excitation shown in panel a. (c). Multiple cycles of photobleaching and photoreversibiliy of mfRFP-A, photobleaching is induced using the 594nm laser channel at 100% power for 120 seconds followed by induction of photo-reversibility with the 488nmnm laser channel. (d) Reference trace of 5 cycles of photobleaching and photoreversibility in 120 frames image of mCardinal and mfRFP-A. (e) (above) Raw image of each cycle of photobleaching/photoreversibility at every 30frame (T1-T5) in a 120frame acquisition, (middle) the overlay of the unmixed result mCardinal-Mito (Magenta) and mfRFP-A-cytoplasm (green) of each cycle, (bellow) a magnified FOV of mito dynamic changes across the timepoints, (T1-T5)

During imaging acquisition, we noticed that, unlike TMI and ExNEEMO, which rely on fast photoswitching FPs^22,23^, the TDM imaging with mCardinal and mfRFP allowed the acquisition of multiple frames before achieving a sufficient level of photobleaching required for reliable unmixing. We took advantage of this feature to test the ability of volumetric TDM imaging of cells, which was not previously performed with TMI or ExNEEMO. For this, we acquired time-lapse z-stack images of HeLa cell co-expressing H2B-mfRFP and keratin-mCardinal fusions using a spinning disk confocal microscope. Then, each z-plane was unmixed independently using the same reference traces (**Figure 3s,t**). The output images demonstrated patterns recapitulating stereotypical structures of the H2B and keratin in mfRFP and mCardinal channels, respectively. The summation of the mfRFP and mCardinal intensity profiles along selected linecuts reconstituted those from raw input images, suggesting TDM crosstalk between two FPs was minimal (**Supplementary Figure 5**). As a result, we obtained a z-stack set, allowing us to render a 3D dual-color image of the cell (**Figure 2u**). In conclusion, mfRFP and mCardinal enabled straightforward 3D multiplexed imaging using a single channel under a conventional confocal microscope.

We successfully achieved reproducible unmixing using various combinations of FPs across different imaging modalities, as described above. However, during image processing, we encountered several challenges that significantly impacted the outcome of TDM unmixing. These challenges should be carefully discussed and presented to inform potential end-users about key experimental considerations when designing TDM experiments. In some cases, we observed noticeable differences in photobleaching rates between brighter and dimmer subpopulations of the same fusion protein. For example, when analyzing mitochondria-targeted mCardinal, we found considerable discrepancies in photobleaching kinetics between mitochondria with higher versus lower fluorescence intensity. This heterogeneity of photobleaching led to a reference trace that did not accurately represent the entire population, resulting in deviations in the unmixing outcomes. Specifically, when we used a reference trace derived from either the brighter or dimmer mitochondria, the resulting unmixing varied considerably (**Supplementary Figure 6**). Additionally, unmixing was compromised when two or more FPs exhibited highly similar photobleaching kinetics. In such cases, the algorithm was unable to effectively separate the signals, leading to incomplete or inaccurate unmixing results (**Supplementary Figure 7**). Therefore, careful selection of FP combinations and validation of photobleaching reference traces are crucial to ensuring reliable TDM analysis. In summary, these challenges highlight the importance of implementing rigorous controls and optimization strategies when designing TDM experiments. Future studies should explore improved approaches for reference trace selection, including adaptive correction methods, to enhance the robustness and reproducibility of unmixing outcomes.

### mfRFP-A enables two-color TDM imaging of subcellular dynamics in live cells

During the recording of the reference photobleaching traces, we noticed that mfRFP-A fluorescence could recover after some time in darkness, while other tested far-red FPs remained photobleached. We hypothesized that this reversibility could be facilitated by blue-shifted illumination, similar to other reversibly photoswitchable RFPs (rsRFPs), such as, for example, rsTagRFP^24^. To test this, we recorded the fluorescence of mfRFP-A under continuous wide-field excitation for 5 minutes, followed by cyan illumination (478/25 nm). mCardinal, mCardinal-A, E2-Crimson, mNeptune, and mfRFP expressed in the cytoplasm of HeLa cells were used as references under identical imaging conditions. Indeed, after almost complete photobleaching, the fluorescence of mfRFP-A could be partially recovered to about 54% of the initial brightness under cyan illumination within 5 minutes (**Figure 3a**). In contrast, all other tested far-red FPs showed either no fluorescence recovery (mfRFP) or very limited photoswitching, with fluorescence increases of 19% for mCardinal, 13% for mCardinal-A, 40% for E2-Crimson, and 29% for mNeptune, compared to 134% for mfRFP-A (**Figure 3b**). Moreover, mfRFP-A could undergo several ON/OFF cycles of photoswitching under 594/478 nm illumination with approximately 2-fold fluorescence contrast (**Figure 3c**). Photoswitching enables repetitive TDM imaging of the same live cell at multiple time points, as demonstrated for green and red rsFPs by Qian et al.^25^ To explore this capability, we used the mfRFP-A/mCardinal pair to visualize mitochondrial dynamics in HeLa cells. Live HeLa cells expressing mfRFP-A in the cytoplasm and mCardinal in mitochondria were imaged over five photobleaching/photoswitching cycles. Each cycle comprised 20 frames acquired in the far-red channel, followed by illumination with 478 nm light to photoswitch mfRFP-A, with a total duration of 1 minute per cycle (**Figure 3d**). After image acquisition, each cycle was processed independently for unmixing, yielding five two-color images recorded at 30-second intervals, enabling mitochondrial dynamics tracking (**Figure 3e**). Altogether, these findings establish mfRFP-A as the first reversibly photoswitchable GFP-like fluorescent protein with a fluorescence emission maximum beyond 650 nm and demonstrate its applicability for time-lapse imaging of subcellular dynamics in live cells using TDM.

### Benchmarking of mfRFP in mammalian cells

Since mfRFP demonstrated high photostability in solution and cultured cells, we decided to systematically benchmark it against other state-of-the-art far-red FPs in live and fixed mammalian cells to assess its broader applicability. For comparison, we selected E2-Crimson, mCardinal, mCarmine, mGarnet, mNeptune, and emiRFP670, which all share very similar spectral profiles, except for emiRFP670, which is more red-shifted compared to all others (**Table 1**). In live HEK cells, the brightness of mfRFP was comparable to that of mCarmine and mGarnet, when quantified by normalization to green fluorescence of EGFP co-expressed via P2A peptide (**Figure 4a**). However, intracellular brightness of E2-Crimson, mCardinal, and mNeptune was 2.6-, 4.7-, and 3.8-fold higher than that of mfRFP (**Figure 4a**). In PFA-fixed HeLa cells, mfRFP was only 2.4-, 2.1-, and 2.9-fold dimer than E2-Crimson, mCardinal, and mNeptune, respectively, indicating higher resistance to fixation compared to mCardinal and mNeptune (**Figure 4b**). Upon continuous wide-field illumination in live cells, mfRFP possessed the highest photostability, followed by mCarmine, mGarnet, mCardinal, mNeptune, and E2-Crimson (**Figure 4c**). In the fixed cells, the photostability trend was different, with mCarmine being the most photostable, followed by emiRFP670, mfRFP, E2-Crimson, mCardinal, and mNeptune (**Figure 4d**). We also observed minor initial photoactivation of mCardinal, mGarnet, and mNeptune in live cells, which, however, was diminished in the fixed cells (**Figure 4d**). In addition, fusions of mfRFP with microtubule-associated end-binding protein 3 (EB3) protein, partial cytochrome C oxidase subunit VIII (COX8A), keratin, and histone 2B (H2B) demonstrated proper localization in cultured HeLa cells (**Figure 4e**). In summary, in cellulo characterization showed that while mfRFP exhibited lower brightness than E2-Crimson, mCardinal, and mNeptune, it outperformed all of them in terms of photostability and resistance to fixation (**Table 1**). These properties make mfRFP a promising fluorescent tag for applications requiring extended imaging and reliable signal retention, particularly under high-intensity illumination.

**Figure 4.**
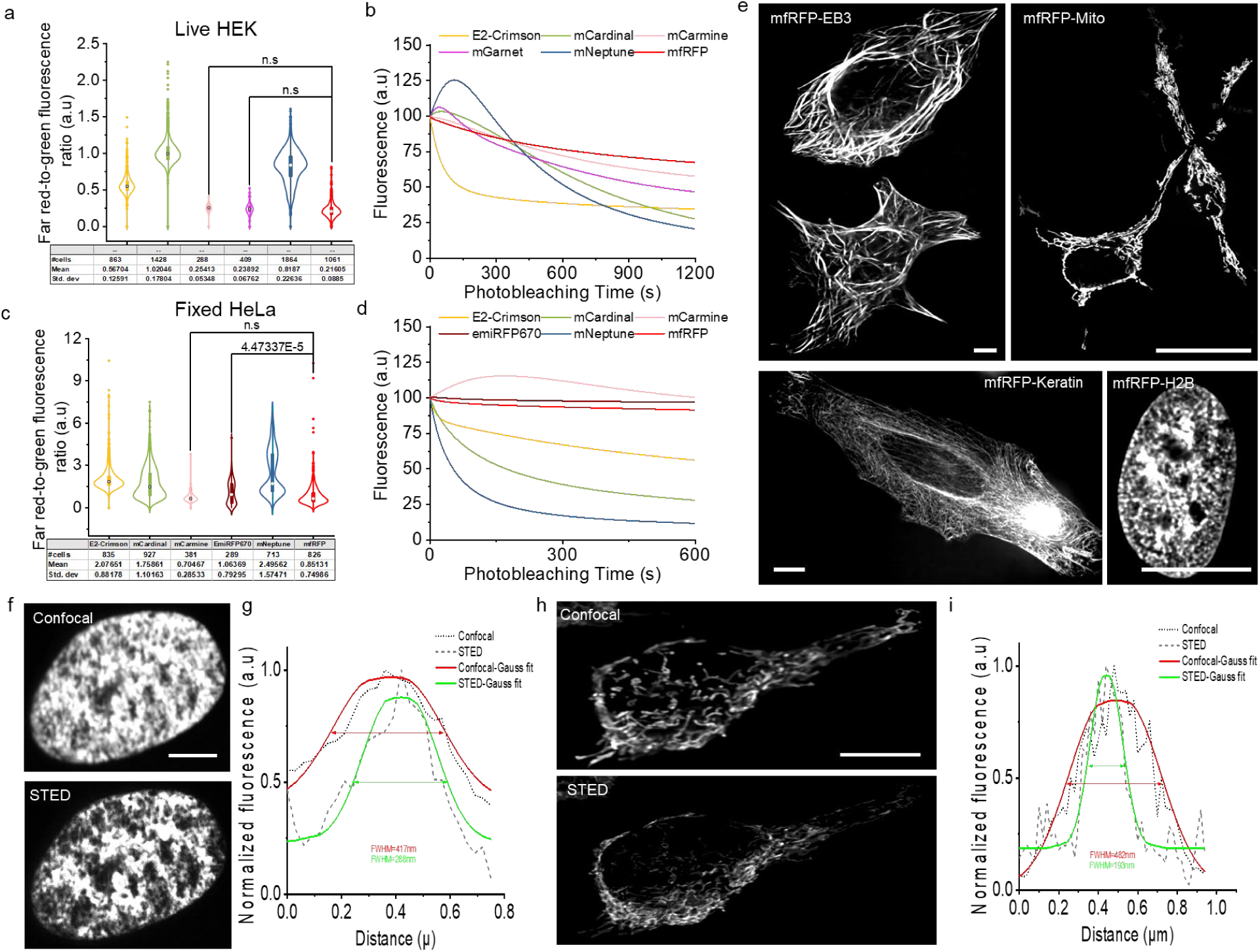
Characterization of mfRFP in mammalian cell culture. (a) Violin plot depicting far red-to-green fluorescence ratio in live HEK cells co-expressing E2-crimson, mCardinal, mCarmine, mGarnet, mNeptune, and mfRFP with EGFP via P2A self-cleaving peptide, 2 independent transfections each; imaging conditions: RFPs excitation 594nm, emission 660nm; GFP: 594nm excitation, emission) boxplot: box range; 25th and 75th percentile, center white dot; median, width density; frequency, black dots; outliers. (b) Normalized photobleaching curve for E2-crimson, mCardinal, mCarmine, mGarnet, mNeptune, and mfRFP expressed in HEK cells, 2 independent transfection each; imaging conditions for HEK cells: 594nm excitation, 660nm emission, power 20%, exposure time 20ms) (c) A Violin plot depicting far red-to-green fluorescence ratio in fixed HeLa cells co-expressing E2-crimson, EmiRFP870, mCardinal, mCarmine, mNeptune, and mfRFP with EGFP via P2A self cleaving peptide, 2 independent transfections each; imaging conditions: RFPs excitation 594nm, emission 660nm; GFP: 594nm excitation, emission) boxplot: box range; 25th and 75th percentile, center white dot; median, width density; frequency, black dots; outliers (d) Normalized photobleaching curve for E2-crimson, EmiRFP870, mCardinal, mCarmine, mNeptune, and mfRFP expressed in HEK cells; 2 independent transfection each; imaging conditions for HEK cells: 594nm excitation, 660nm emission, power 20%, exposure time 20 ms(c) (e) Representative images of mfRFP in various fusions (clockwise from top left) EB3-mfRFP, Mito-mfRFP, Keratine-mfRFP and H2B-mfRFP (n>5 cells for each construct from two independent transfections). Scale bar 25μm EB3,CytERM,Mito and H2B were obtained under identical microscopic setup Nikon Confocal 100X 500us exposure time at 594nm laser channel while Keratine was obtained with Nikon Confocal 500us exposure time at 594nmSoRa laser channel and Mito was obtained using the Olympus Confocal microscope 60X. (f) Confocal (top) and STED (bottom) images of H2B-mfRFP fusion. Scale bar 25 um. (g) Cross sections of the nucleus marked in orange line in (f), the line profile of the confocal (black, dot) and STED (dash, gray) image obtained from the solid orange line, the solid lines (red, confocal) (green, STED) represent Gaussian fits used to determine the full width at half-maximum (FWHM) values. (h) Confocal (top) and STED (bottom) images of Mito-mfRFP fusion. Scale bar 25 um. (i) Cross sections of the nucleus marked in orange line in (f), the line profile of the confocal (black, dot) and STED (dash, gray) image obtained from the solid orange line, the solid lines (red, confocal) (green, STED) represent Gaussian fits used to determine the full width at half-maximum (FWHM) values

Due to its relatively high intrinsic photostability, we evaluated the compatibility of mfRFP with STED super-resolution microscopy, which operates at higher excitation powers than conventional diffraction-limited microscopy and, therefore, requires more photostable FPs. Using a commercially available STED system, we were able to acquire super-resolution imaging of H2B and COX8A fusions achieving 40 nm resolution for mitochondrion (**Figure 4f-i**), comparable to that for mCardinal and miRFP670 (**Supplementary Figure 8**). For reference, the mfRFP photobleaching rate under STED was comparable to that for mCardinal and miRFP670, which are state-of-the-art FPs for STED imaging (**Supplementary Figure 9**). However, mfRFP exhibited slightly higher photostability over mCardinal and miRFP670 when compared under point scanning confocal excitation (**Supplementary Figure 9**). These findings support mfRFP as an FP tag suitable for STED microscopy, offering comparable photostability relative to other far-red FPs in both standard and high-intensity imaging conditions.

### mfRFP expression in model organisms in vivo

Red-shifted fluorescence is particularly beneficial for imaging mammalian tissue and small model organisms such as *C. elegans* and zebrafish due to reduced autofluorescence, low light scattering, and minimal absorbance of tissue at longer wavelengths of the visible spectrum^26^ From many of our previous studies on FPs, we noted that the performance of FPs in cultured cells, such as HeLa and HEK, often did not translate well into performance in neurons in vivo^27–31^ Particularly, reduced intracellular brightness and poor neuronal labeling, manifested as intracellular aggregation and limited trafficking to distal neurites, are among the most common drawbacks of FPs when expressed in neurons in vivo^32^. Given these common limitations, we sought to evaluate the utility of mfRFP for neuroimaging in model organisms, including mice, zebrafish larvae, and *C. elegans*. We started by co-expressing mfRFP with EGFP in the mouse cortex through neonatal injection of the rAAV2/9-CAG-mfRFP-P2A-EGFP virus, followed by histological analysis of the targeted brain regions in four-week-old mice using high-resolution imaging. Interestingly, while both neurons and astrocytes were visible in the green channel, only astrocytes appeared in the far-red channel. This observation suggests that mfRFP may exhibit either low expression levels or reduced intracellular brightness in murine neurons (**Figure 5a**). Next, mfRFP was transiently co-expressed with mClover3 under a pan-neuronal promoter in zebrafish larvae and visualized at 4 days post-fertilization (**Figure 5b,c**). High-resolution confocal imaging revealed that mfRFP was evenly distributed throughout the neuronal cytoplasm without visible aggregation, including in the hindbrain and spinal cord, compared to that of mCardinal (**Fig. 5c; Supplementary Figure 10**). Lastly, we demonstrated that mfRFP can be used for neuronal labeling in *C. elegans* without forming aggregates. However, visualization of small neuronal processes was less efficient compared to mNeonGreen, as observed in the image overlay (**Figure 5d**). Overall, mfRFP was suitable for neuroimaging of model organisms.

**Figure 5.**
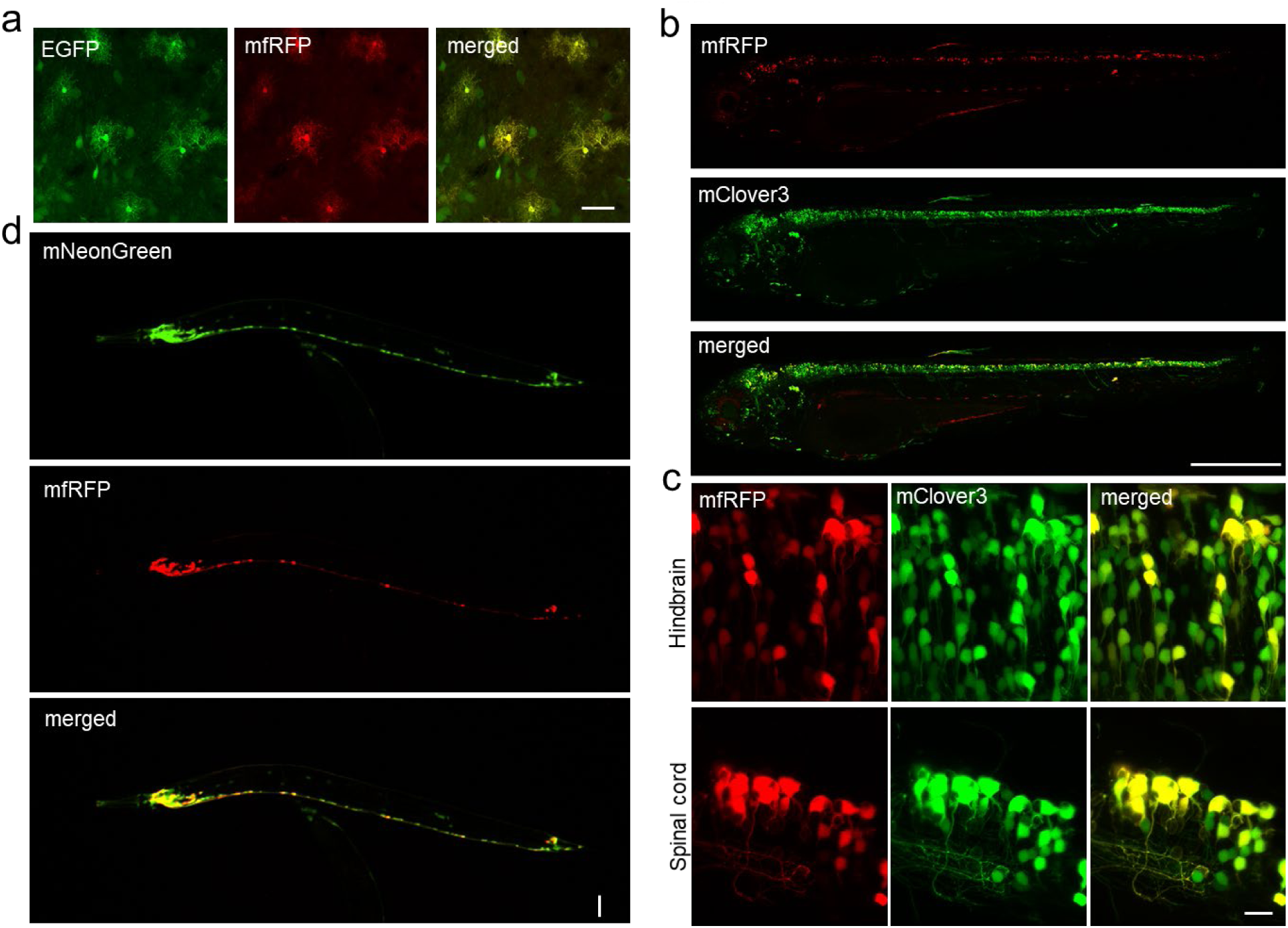
(a) Representative overview images of 4dpf zebrafish showing mfRFP (upper, red), mClover3 (middled green), and merged fluorescence (lower) throughout the brain and spinal cord. Green channel: excitation 488 nm Argon laser, emission 496-530 nm; Cy5 channel: excitation HeNe 633 nm laser, emission 645-780 nm. Scale bar: 500 µm. (b) Higher magnified images of hindbrain (above) and spinal cord (below) neurons showing mfRFP (red), mClover3 (green), and merged fluorescence in larval zebrafish at 4 dpf (right). Green channel: excitation 488 nm Argon laser, emission 496-530 nm; Cy5 channel: excitation HeNe 633 nm laser, emission 645-780 nm. Scale bar: 15 µm. (c) From right to left, representative image of mice brain hippocampus region expressing mfRFP (red), EGFP (green) and merged(yellow). Green channel Excitation: 488nm mfRFP channel Excitation: 594nm. Scale bar 25µm. (d) Representative whole-body image of *C. elegans* co-expressing mNeonGreen (green) and mfRFP (red) under pan-neuronal promoter tag168. Green channel excitation: 488 nm; mfRFP channel excitation: 594 nm. Scale bar 25 µm.

## DISCUSSION

In this study, we present a set of far-red FPs that have been specifically engineered to exhibit distinct photobleaching kinetics. Our motivation for developing these FPs was to explore their potential for TDM imaging, leveraging differences in photostability rather than relying on reversible photoswitching, which is employed in unmixing techniques such as TMI^11^ and ExNEEMO^10^. We suggested that FPs with primarily irreversible photobleaching characteristics could also be effectively employed for TDM. Our findings confirm that such FPs can indeed be successfully unmixed in the temporal domain across multiple imaging modalities, including widefield, spinning-disk confocal, and point-scanning confocal microscopy. Notably, because reversible switching is not required, this approach allows TDM to be extended to imaging systems that do not support reversibly photoswitchable FPs, such as point-scanning confocal microscopes, thus broadening the accessibility of this technique. Furthermore, it requires only 20 to 40 frames to achieve unmixing for up to 3 FPs with distinct photobleaching kinetics, and the image processing can be executed on a standard computer using MATLAB software.

While we successfully unmixed several FPs, we also identified specific challenges, as presented and described in detail in the Results section. One of the key technical issues arises from the fact that within the same cell, different subcellular structures expressing the same fusion protein may exhibit slightly different photobleaching kinetics, likely due to variations in brightness or the local environment. In such cases, unmixing becomes suboptimal. Therefore, end users should carefully select combinations of FPs for TDM and validate them in desired fusions under conditions tailored to the specific requirements of the intended application. An additional experimental consideration is to match the brightness of the used FP fusions, as this may enhance unmixing and reduce cross-bleeding of the brighter FPs into the channel of the dimmer FP. Once the combination of FP fusions is selected, it might also be suggested to perform validation of the TDM unmixing with FLIM or immunostaining for a subset of cells. Interpretation of the unmixing results should also take into account the potential limitations of the technique mentioned above. However, it is important to note that these challenges are not unique to the TDM algorithm described here but are also prevalent in the TMI and ExNEEMO methods, as they all share the same underlying mathematical principles. Moreover, FLIM unmixing also exhibits similar limitations, which are associated with an insufficient difference in fluorescence lifetime (as shown in **Supplementary Figure 7**) and environmental sensitivity of fluorescence lifetime of FPs, even within the same cells^33,34^ in addition to the high-temperature sensitivity of fluorescence lifetime^35^ While sharing common limitations, the TDM approach offers specific advantages over TMI and ExNEEMO, making it more practical for FP unmixing. First of all, TDM can be applied to any spectral class of common FPs, which are now more than 1,000 according to fpbase.org, as long as they exhibit distinct photobleaching kinetics. While TMI and ExNEEMO require reversibly photoswitchable FPs, which are now limited to a handful of green and red rsFPs. Second, the utilization of rsFPs relies on specific imaging acquisition protocols with fast switching between excitation and photoswitching illumination. This image acquisition procedure can be performed only with fully automated and raid imaging setups, increasing requirements for microscopy hardware. In contrast, TDM datasets are acquired in a single channel without a need for additional illumination or automated control of the hardware (rather than just the camera itself). This feature makes TDM more practical and extends its applicability to point-scanning confocal microscopy. At the same time, it enables 3D imaging with confocal microscopy, which was not realized with TMI and ExNEEMO. Overall, TDM offers a more accessible alternative to FLIM, TMI, and ExNEEMO, with minimal requirements for the imaging hardware and providing broader coverage of the spectral class of the utilized FPs.

In addition to the new, immediate applications TDM offers for cell biologists and microscopists, it also opens up interesting opportunities for protein engineers. For decades, protein engineers have focused on optimizing the FPs to develop the most photostable variants. Consequently, high photostability has traditionally been a major selection criterion. However, emerging techniques like TDM now create a demand for spectrally similar FPs with diverse photobleaching properties, providing protein engineers with a broader range of selection criteria when evolving new FPs. We expect that this demand will promote further generation of novel FPs in all spectral classes. The TDM technology can be also improved on the image analysis aspect. It might be possible to improve existing unmixing algorithms to account for spatially variable photobleaching rates. For example, future algorithms could incorporate reference traces specific to different brightness levels, enhancing the robustness of TDM in heterogeneous cellular environments. Altogether, we believe that TDM holds great potential for multiplexed imaging, and the technology can be enhanced with new sets of optimized FP combinations and more powerful unmixing algorithms.

## METHOD SECTION

### Molecular cloning and mutagenesis

The mCarmine,^16^ mMaroon,^17^ mGarnet2 (ref.^17^) genes were synthesized *de novo* by Synbiob Gene Technology Co., Ltd., (China), based on the sequences reported in the original publications, and cloned into pHybrid-Electra1 (swapping Electra1 gene; WeKwikGene plasmid #0000202) and pAAV-CAG-mCardinal-P2A-GFP (swapping mCardinal gene; WeKwikGene plasmid #0000002) plasmids. The mCardinal and TagRFP658 genes were PCR amplified from pAAV-CAG-mCardinal-P2A-GFP (WeKwikGene plasmid #0000002) and pAAV-CAG-TagRFP658-P2A-GFP (WeKwikGene plasmid #0000244) plasmids, respectively. For expression in mammalian cell lines, the target plasmids were cloned in two steps. First, EBFP2 in pAAV-CAG-EBFP2-P2A-GFP (Addgene plasmids #184938) was swapped with the gene of the selected.

Synthetic DNA oligonucleotides used for cloning were synthesized by Tsingke Biotechnology (China). PrimeStar Max master mix (Clontech) was used for high-fidelity PCR amplifications. Restriction endonucleases were purchased from (New England Biolabs, United States) and used according to the manufacturer’s protocols. DNA ligations were performed using T4 DNA ligase (New England BioLabs) or the NovoRec plus one-step cloning kit (Novoprotein). Small-scale isolation of plasmids was done with the QIAGEN plasmid mini kit (Germany). Sequencing was performed by the Sanger method (Zhejiang Youkang Biological Technology, China).

Error-prone PCR of the TagRFP658 gene was performed using Mutazyme II DNA polymerase (Agilent, United States) under high mutation rate conditions (9–16 mutations per kilobase pair) and subcloned into the pBAD-HisD-Electra1 plasmid (WeKwikGene plasmid #0000248). The generated libraries were electroporated in TOP10 cells (Tsingke, China) and expressed in LB media supplemented with ampicillin (Amp+: 100 mg/μl) and 0.02% L-arabinose (Ara) for 24 h at 37°C. For quality control, after electroporation an aliquot (1 µl) of each library was plated on LB/agar plates to estimate the library size and confirm ligation efficiency and mutation rate by sequencing 20 randomly picked colonies. All generated libraries contained 2–70×10^6^ independent clones with six to eight nucleotide mutations per gene. Bacterial cultures expressing the libraries were sorted using FACS (BD FACS Melody, United States) with 640 nm excitation light. The collected cells were plated on LB/agar plates supplemented with Amp+ and Ara and incubated overnight. The colonies were imaged in a green channel: (Ex: 580-600 nm; Em: 610LP) using an SZX16 fluorescence stereomicroscope (Olympus, Japan) equipped with Spectra III Light Engine (LumenCor, United States) and cooling color digital camera (BGIMAGING, China). The selected colonies were cultured in LB medium containing Amp+ and Ara in 24-well deep well plates for 24 h at 37°C and 16-18h at 18°C, treated with B-PER to extract proteins (Thermo Fisher, United States or Shenggong, China), and analyzed using a fluorescent plate reader (Varioskan LUX, Thermo Fisher Scientific, United States). Photostability of the 10-20 brightest clones were measured in droplets in mineral oil under continuous wide-field illumination using Nikon Ti2-E wide-field fluorescence microscope equipped with a Spectra III Light Engine (LumenCore, United States), 680 nm CNI laser, and an ORCA-Flash 4.0 V3 sCMOS camera (Hamamatsu) with ×10 0.45 numerical aperture (NA) and ×20 0.75NA objective lenses (Nikon) controlled by NIS Elements AR 5.21.00 (Nikon).

### Protein purification and in vitro characterization

For protein purification, TagRFP658, mGarnet2, NIRFP1, mMaroon, mCardinal, mCardinal-A, mfRFP, mfRFP-A genes cloned into the pHybrid vector were transformed into TOP10 *E. coli* strain (Biomed), the bacterial cells were grown in RM medium supplemented with 100 μg/mL ampicillin and 0.02% w/v L-rhamnose (Sangon Biotech (Shanghai) Co., Ltd.) at 37°C for 24 h followed by incubation at 18°C in shaking incubator at 220 rpm. The products were centrifuged at 4000 g at 4°C for 15 min. The cell pellets were resuspended in PBS buffer, pH 7.4, supplemented with 300 mM NaCl, and lysed by sonication on ice. Proteins were purified utilizing metal affinity chromatography using Ni-NTA resin (20503ES50, Yeasen Biotechnology) according to the manufacturer’s instructions. The proteins were eluted with PBS containing 400 mM imidazole, and 300 mM NaCl in 1x PBS at pH=8.0, followed by dialysis overnight against PBS buffer, pH 7.4. To determine the absorption spectra of the purified FPs, an absorbance scan (260-650nm) was measured with the UV-vis absorption spectra machine using UV-VIS-NIR UV3600Plus spectrophotometer (Shimadzu, Japan). The steady-state fluorescence spectral peaks were obtained using a cuvette filled with 100 μl of the eluted protein measured using a Fluorolog3 spectrofluorometer (Jobin Yvon, Horiba, United States).

The extinction coefficients were measured by alkali-denaturation in 1M NaOH to determine the mature chromophore concentrations, assuming that the extinction coefficient of chromophores is 44,000 M^−1^cm^−1^ at 447 nm in 1M NaOH. Briefly, 150 μl of the protein was diluted to a peak absorbance value between 0.5 and 1.0, measured from 200 nm – 800 nm, followed by mixing 1:1 with 150 μl of 2M NaOH and remeasuring absorbance. The absolute fluorescence quantum yield (QY) was measured using an FLS1000 spectrometer equipped with the integrating sphere accessory (Edinburgh Instruments, UK) according to the manufacturer’s protocol. Briefly, 2– 10 ul of the eluted protein was diluted to an Optic density of 0.1-0.5 to minimize the inner filter effect, and the proteins were excited at 580 nm and measured by the ratio of the integrated fluorescence values (in the range of 590–800 nm) in PBS buffer and that of the scattering values, FluOracle Software (Edinburgh Scientific, UK) was used for computing the resulting quantum yields value.

### Mammalian Cell Culture

HeLa cells (ATCC CCL-2) and HEK293FT cells (Invitrogen) were grown and cultured in Dulbecco’s Modified Eagle Medium (DMEM) (Servicebe DMEM/High Glucose) supplemented with 10% fetal bovine serum and 1% penicillin/Streptomycin (PS) incubated in 37°C and 5% CO2 humidified incubator machine. Transient transfections were performed in 24-well plates (NEXT, Lot: 040222BH01-0422B) or 35mm MatTek dishes (MatTek Life Sciences, P35G-1.5-14-C) coated with Matrigel (BD Biosciences, 356235). Cells were transiently transfected at 40-60% confluency using Hieff Trans™ Universal Transfection Reagent (Yeasen, China) according to the manufacturer’s instructions.

Cells in 24 well plates were initially washed twice with PBS and fixed with 4% paraformaldehyde (PFA) at room temperature for 10 min, The fixed cells were washed twice with PBS and subsequently mounted with Prolong Diamond Antifade mountant (Invitrogen Thermofisher scientific) on microscope slides CITOTEST 25*75mm, 1mm-1.2mm, the 24 well plates and dishes were left in the dark 12-24 hours for curing.

### Fluorescence microscopy

Intracellular brightness and photostability measurements were measured with the Nikon Ti2-E widefield microscope equipped with Spectra III Light Engine (LumenCore), the ORCA-Flash 4.0 V3 sCMOS camera (Hamamatsu), and 20x/0.75 objective lens controlled by NIS Elements software (NIS-Elements AR 5.30.00 64-bit) using, GFP (excitation 488/28 nm, emission 535/46 nm), mfRFP (excitation 594/23 nm, emission 610LP) channel.

To quantify intracellular brightness in HeLa cells, cells were imaged in the FITC (excitation 475/28 nm at 5.30 mW/mm^2^ from SpectraIII LumenCor; emission 535/46 nm), and mfRFP (excitation: 594/23 nm at 56–58 mW/mm^2^ from 680 nm CNI laser; emission 610LP) channels. To obtain statistically significant datasets, we performed three independent transfections and analyzed more than 100 regions of interest (ROIs) from each transfection (at least 300 ROIs for analysis in total per construct). ROIs were determined using the auto-detect function of NIS Elements software and limiting the smallest ROI to 50 μm^2^ (the average size of HeLa cell). Mean fluorescence intensities in green (FITC) and mfRFP channels were measured for each ROI, and background fluorescence was subtracted from both channels. The mfRFP-to-green fluorescence ratio was calculated for each construct to compare intracellular brightness under standardized imaging conditions. Dead and shrunk cells and samples with bacterial or yeast contamination were excluded from the analysis.

The photostability of proteins was assessed in live and fixed HeLa cells (transfected as previously described) under continuous (excitation 594nm/23nm, emission 610LP) laser excitation via wide-field microscopy. Photobleaching curves were generated individually for each cell, and the average photobleaching half-life (with standard deviation) was calculated for each protein. Photobleaching experiments were conducted until a 50% fluorescence intensity drop was achieved, where feasible, and the mean photobleaching curve for each protein was reported (averaged across all individual curves). Cells that detached from the substrate or exhibited signs of viability loss during imaging were excluded from data analysis.

Photo-switching experiments were conducted on the same microscope as above, intrinsic photo-switching behaviors of far-red FPs, mfRFP, mfRFP-A, mCardinal, mCardinal-A, mNeptune, and E2-Crimson, were measured under identical imaging parameters and conditions, the transfected cells were imaged using continuous illumination (594 nm laser channel 100% Power, Exposure time 10 ms) for 5 minutes duration followed by 10 minutes period of intermittent blue light pulse (488nm laser channel 20% power, Exposure time 10 ms) at every 5 seconds interval to switch the FPs from the OFF state to the ON state.

### Fluorescence lifetime imaging

Fluorescence Lifetime images were acquired with the Stelaris Falcon 8 FLIM microscope system (Leica, Germany). The laser light was focused using the 40x 0.94 Na (water) objective lens. A dichroic filter at 561nm was employed to separate the excitation and emission light. The Power HyDx photomultiplier detector was employed for the fast lifetime imaging and photon counting. FLIM data acquisitions were analyzed with LAS X FLIM/FCS analysis software (Leica Microsystems, CMS GmbH). Fluorescence lifetime image acquisition settings, 4-line repetitions, and 3 frame repetitions to achieve optimal photon counts, scan speed, 0.097/s frame rate, and pixel size,1024 * 1024, pixel dwell time 1.41us. For TDM application, (x,y,z,t) images were acquired with the confocal microscope setup (594nm excitation at 100% laser power) for each image, after maximum intensity z projection, 19 frames of time-lapse images were obtained for 10 min.

Lifetime analysis and channel unmixing were performed using Leica Application Suite LAS X FLIM/FSC Analysis Software (Leica Microsystems CMS GmbH, version 4.7.0). Multiplexed FLIM images were analyzed via phasor plot analysis. This one-to-one mapping allows for interactive visualization: selecting a specific region (i.e., a cluster of points) on the phasor plot instantly identifies the spatial distribution of pixels in the FLIM image corresponding to that lifetime signature. This approach enables the clear localization and differentiation of distinct fluorophore populations with unique lifetime values. The intensity or color gradient at any point on the phasor plot quantitatively reflects the pixel density (i.e., the number of pixels) sharing identical lifetime characteristics, thereby providing a spatially resolved and quantitative representation of fluorophore dynamics.

### TDM data processing

Time-series fluorescence imaging data were analyzed using Time Domain Multiplexing to decompose mixed fluorescence signals into distinct components based on reference photobleaching profiles. The workflow integrated raw image preprocessing, computational decomposition, and validation using MATLAB (MathWorks R2020b), Nikon NIS-Elements AR (version 5.30.00 64-bit), and Leica STELLARIS FLIM microscopy.

Image denoising were achieved using the inbuilt NIS.ai function a deep-learning-based algorithm within NIS-Elements AR Analysis (5.30.00 64-bit) to reduce background noise while preserving signal integrity Pixelwise decomposition was achieved utilizing the least square regression strategy: for each pixel, the algorithm fits its intensity trajectory across frames to the reference photobleaching curves using the MATLAB least square regression fit function. The optimization strategy computes the optimal coefficient values representing the strength of the match between the pixel’s intensity profile and each reference curve were calculated via ***fmincon*** function (MATLAB’s constrained optimization function) to minimize residuals between the observed pixel trajectory and reference profiles.

Coefficient values representing the strength of the match between the pixel’s intensity profile and each reference curve were normalized to a 0-1 scale; where 0 represents pixels not matching a reference profile, and 1 represents the pixel matching a reference profile with high fidelity.

Normalized coefficients were multiplied by the first frame of the raw image to generate channel-specific images corresponding to each reference profile, isolating fluorescence signals of interest in each channel. The obtained brief movies were analyzed using the Nikon NIS element Advance Research (AR) and Fiji ImageJ software.

The modified TDM code (TMI^11^) was implemented in MATLAB (MathWorks R2020b) to decompose the denoised images into respective channels with improved functionality. Three new folders are created, the code folder, containing the code, its function and the user’s reference profile. The data folder: containing the raw-data file to analyzed. The result folder stores the output image are automatically saved into respective decomposition channels.

Data recognition have been improved from the previous version; image size and dimensions are auto-recognized once uploaded to the data folder facilitating ease of access. In addition, the generated results are in the form of a time-series image, with each frame representing each time frame of the input image.

### Unmixing result analysis

The output images from the Time-Domain Multiphoton (TDM) decomposition algorithm are analyzed for drift (misalignment) and aberrations (distortion) using the Longitudinal fluorescence intensity line profile tool in ImageJ software. To facilitate a direct comparison, line profiles are drawn at identical positions across both the raw images and the decomposition images.

A statistical analysis is then performed by computing the correlation coefficient between the raw image and the merged decomposition images. This assesses the similarity between the original images and the decomposed images, enabling the evaluation of any potential drift or structural distortion/ aberrations introduced by the TDM decomposition algorithm

By using the correlation coefficient as a metric, the degree of similarity between the raw image and the merged decomposition images can be quantitatively determined. This hybrid approach-combining line profile analysis for spatial shifts and statistical metrics for structural fidelity provides a robust method for validating the accuracy of the TDM decomposition algorithm and identifying any potential sources of error.

### Crosstalk measurement

Crosstalk between fluorescence channels was quantified using imageJ software, leveraging the ROI manager tool to systematically analyze signal contributions from fluorescent proteins (FPs) in dual-channel imaging.

Channel 1(target channel) was analyzed by measuring the residual intensity contribution of FPs channel 2(non-target channel) across multiple regions of interest (ROIs). The mean residual pixel intensity of Channel 2 signals in Channel 1 ROIs was calculated and normalized to account for bleed-through. The analysis was repeated with Channel 2 designated as the target and Channel 1 as the non-target to assess reciprocal crosstalk.

Crosstalk values (e.g., bleed-through percentages) were plotted to compare channel-specific interference. This bidirectional approach ensured comprehensive quantification of spectral overlap and cross-channel contamination, enabling precise quantification of fluorescence bleed-through.

### STED microscopy

STED imaging was obtained at the imaging facility of The Institute of Life Science, Zhejiang University, Hangzhou. Images were acquired with the Abberior Instruments, Facility Line, Germany, coupled with Olympus Microscope controlled by Lightbox, Imspector software equipped with a 60× 1.4 oil immersion lens. All images were excited with the Abberior Star Red dye (630-650nm) excitation laser and 775nm STED depletion laser.

For typical STED versus confocal images acquisition, the pixel size for STED and confocal images are identical for each acquired image (30nm-45nm), the pixel dwell time for STED images was set at 30 to 45 µs, and for the confocal images, 5 – 35 depending on the observed photon count, the relative laser intensity for Confocal image excitation laser was set at 5%-70% depending on the observed photon count and STED excitation laser 5%-70% and STED 775 nm depletion laser 5% - 20% were applied.

For time series image acquisition, the relative excitation laser intensity for all STED images was 5% and 70% for confocal depletion, accumulation, pixel dwell time, and pixel size were set at 1, 40 µs and 30nm, respectively for all acquisition, Data were collected for 30 iterations with 1-second interval for STED images and 200 iterations and 1s interval for confocal images.

### Image processing and analysis

Image processing such as z-stacks projection of acquired z-stack series, for maximal intensity projection (image>Stacks>Z-project>Maximum intensity projection) Drift correction was corrected using Image registration plugin prompt was used to align time-series images with shift for correction (Plugin>Registration>Linear Stack Alignment with SIFT), The duplicate function was used to truncate time series image frames (image>duplicate)

ImageJ’s built-in function, ROI manager (Analyze>Tools>ROI manager), was used to obtain the time-lapse data, and the Photon count data were obtained by plotting the stack profile.

To calculate the full width at half maximum (FWHM), the regions of interest were highlighted with the ImageJ built-in line tool, set to a width of 3-pixel average for all data measured. The acquired line profiles are plotted with the OriginPro Software and both confocal and STED data are fitted with Gaussian curve for peak analysis to obtain their full width at half maximum (FWHM).

### Animal procedures and experiments

All animal maintenance and experimental procedures for mice were conducted according to the Westlake University animal care guidelines, and all animal studies were approved by the Institutional Animal Care and Use Committee of Westlake University under animal protocol no. 19-044-KP-2. Procedures involving experimental animals are reported in accordance with Animal Research: Reporting of In Vivo Experiments (ARRIVE). Mice were maintained at strict barrier facilities with macroenvironmental temperature and humidity ranges of 20–26 °C and 40–70%, respectively. Food and water were provided ad libitum. The rooms had a 12-hour light-dark cycle. The housing conditions were closely monitored and controlled. C57BL/6 mice supplied by the Animal Facility at Westlake University were used without regard for sex.

All the viruses were purchased from Shanghai Sunbio Medical Biotechnology Co., Ltd (China). For protein expression across the cortex, viruses (∼ 10^12^ GC/ml) were injected pan-cortically into P0 pups with a Hamilton microliter syringe. To induce hypothermic anesthesia, the pups were put on ice for 5 min until they stopped responding to gentle squeezing on the limbs. For each hemisphere, 0.5 μl of virus solution supplemented with 10% FastGreen dye (SigmaAldrich) was injected under the skull and above the dura manually. After injections on both hemispheres, the pups were moved onto a heating pad maintaining 37°C for a 5-minutes recovery. After the pups regained responses to gentle squeezing, they were returned to the home cages.

For brain slice preparation, the 6–8-week-old mice, regardless of sex, were anesthetized with phenobarbital via intraperitoneal injection; after the surgical plane was reached, intracardiac perfusion was performed at room temperature with 5 ml PBS buffer followed by 50 ml of 4% PFA solution. Decapitation was swiftly followed by removing the skin from the head, exposing the cranium; brain extraction was performed by a rostral midline cut to remove the cranium, exposing the left and right hemispheres; then, the brain was gently peeled from the base of the skull, stored in 4% PFA for approximately 16 hours. The following day, the brain was washed in PBS, followed by a coronal dissection of the fluorescence region (mid-brain) identified with the aid of a lens. The region of interest was immediately glued on a cutting stage of a vibratome (Leica Vt 1200S vibratome). The parameters of the vibratome were optimized for slicing at room temperature (thickness=50 µm) the slices were maintained in PBS at room temperature and mounted on an imaging slide with a Fluoromount mounting medium for confocal imaging.

### Expression of FPs in neurons of C. elegans

For expression in *C. elegans*, the target genes were codon-optimized using the *C. elegans* codon adapter application (https://worm.mpi-cbg.de/codons/cgi-bin/optimize.py) with the insertion of one intron and de novo synthesized by Tsingke Biotechnology Co., Ltd, China). The optimized genes were cloned into pSF11 vector (WeKwikGene plasmid #0000107) under pan-neuronal promoter regulation tag-168. The transgenic lines were prepared by SunyBiotech Co. Ltd (China) according to standard protocols. Briefly, wild-type N2 worms were co-injected with two plasmids pSF11-wmfRFP and pSF11-wmNeonGreen (WeKwikGene plasmid #0000100) with final concentrations 10 ng/μl each. wmNeonGreen was used as a selection marker expressed in most somatic cells. Transgenic lines were selected by green fluorescence and confirmed by sequencing the target transgenes. All worm lines were maintained and selected under the GFP channel regardless of far-red fluorescence until F4. The L4 hermaphrodite adults were transferred on glass slides precoated with 5% agar pads according to previously reported protocol (www.wormatlas.org/agarpad.htm) with a minor modification: 25 mM levamisole (lot#A506644, Sangon Biotech Co., Ltd, China) in M9 was used instead of sodium azide (NaN3) to immobilize worms. Structural imaging was performed under Expression of FPs in neurons of living zebrafish larvae

The zebrafish experiments were carried out according to EU guidelines and German legislation (EU Directive 2010_63, license AZ 325.1.53/56.1-TU-BS). The DNA fragment carrying 10xUASE1b promoter fused to mfRFP/mCardinal-P2A-mClover3-SV40pA was inserted into the Gal4 plasmid, pTol2-nbt3002n: KalNFB-3xmir1T-syt2a (Addgene 196900), generating a Gal4/UAS-mediated expression construct inducing co-expression of NIR FP and mClover3 in neurons throughout the zebrafish brain and spinal cord. These plasmids were injected into pigmentation compromised brass embryos at 1-2 cell stages together with Tol2 mRNA (1.5 nl of injection mix containing 25 ng/μl of both pTol2-plasmid, and Tol2 mRNA), which allowed for efficient chromosomal integration of transgenes, resulting in neuron specific expression in a semi-mosaic manner in zebrafish larvae. 4 days post fertilization (dpf), positive zebrafish larvae were anesthetized with 0.02% Tricaine (E10521, Sigma-Aldrich) and embedded in 1.0% low-melting agarose (A9414, Sigma-Aldrich), followed by being subjected to fluorescence imaging using laser scanning confocal microscope (TCS SP8, Leica Microsystems, Wetzlar Germany) with 40 × NA1.1 water objective. Overview images of zebrafish showing mf-RFP, or mCardinal fluorescence, were acquired (excitation: 633 nm from a HeNe laser, emission 645-780 nm) using the Tile scan mode, and assembled into tiled images using LAS X software (Leica Microsystems, Wetzlar, Germany). For two-color imaging, neurons expressing each NIR FP (excitation: 633 nm from a HeNe laser, emission 645-780 nm) and mClover3 (excitation: 488 nm from an argon laser; emission 496-530 nm) in the hindbrain and spinal cord were imaged simultaneously. Acquired images were processed with FIJI to measure the fluorescent intensity ratio of each NIR FP to mClover3 in each neuron. The assessment of in vivo photostability of each NIR FP fluorescence were conducted with spinal cord neurons from 4 dpf larval zebrafish (40 neurons from 4 independent zebrafish) using laser scanning confocal microscope (excitation: 633 nm from a HeNe laser, emission 645-780 nm) with 40 × NA1.1 water objective (TCS SP8, Leica). A single plain image (optical section: 1.271 μm, image size: 96.88x96.88 μm) encompassing 5-10 fluorescent positive neurons was continuously recorded for 3 min with 70% laser power in the software setting. Acquired images were processed with Fiji, and the fluorescent intensity in individual neurons for each time point was measured.

## Data availability

The in vitro and in vivo measurements and. raw data, including raw images essential to the work, are available online as source data files and provided in the Supplementary Information. The complete datasets, including raw images, are available at FigShare (https://10.6084/m9.figshare.25853149).

## Code availability

Code is available under the BSD-2-Clause license on GitHub (https://github.com/Fakorede-Olumayowa/Temporal_Domain_Multiplexing.git) and Gitlab (https://gitlab.wllsb.edu.cn/blank22/kiryl-piatkevich/-/tree/master/Alex/Temporal%20domain%20multiplexing/Results).

## Acknowledgments

We thank Stavrini Papadaki from Westlake Laboratory for verifying all plasmid sequences and depositing them to WeKwikGene. We thank Hanbin Zhang for help with molecular cloning. We thank the imaging core facility of Zhenzhiang Medical University for providing access to the Abberior STED Facility line microscope system. We also extend our solemn gratitude to Danyu Gu and Dr. Zhong Chen of the physics department laboratory at Westlake University, their instructions and guidance during measurement and data acquisition were highly valued. This work was supported by start-up funding from the Foundation of Westlake University, National Natural Science Foundation of China grant 32050410298 and 32171093 to K.D.P. and by the Ministry of Science and Higher Education of the Russian Federation for the development of the Kurchatov Center for Genome Research 075-15-2019-1659 within the state assignment of NRC "Kurchatov Institute" and by the Russian Science Foundation project 21-74-20135 (protein purification) to F.V.S.

## Author Contribution

K.D.P. and F.V.S. developed mfRFP. K.D.P. and F.O. developed mfRFP-A and mCardinal-A. O.F and J.C. characterized proteins in vitro. O.F. and Z.R. established TDM unmixing procedure. O.F. characterized proteins in cell culture and performed TDM and FLIM analysis. R.W. and O.F. cloned plasmids. X.Y., K.N., and R. W. K. performed experiments in zebrafish and corresponding data analysis. S.P. and O.F. performed *C. elegans* experiments. F.O. performed data analysis and initial interpretation. K.D.P. and O.F., with help from K.N. and X.W., wrote the manuscript.

K.D.P. supervised all aspects of the project. All authors reviewed the manuscript.

## Competing Interest

The authors declare no competing interests.

## Data and materials availability

All data that support the findings of this study are available from the corresponding author upon reasonable request. All sequences of the genes created will be submitted to WeKwikGene, and all plasmids constructed will be submitted to Addgene and WeKwikGene for free distribution.

## Notes

https://10.6084/m9.figshare.25853149

https://github.com/Fakorede-Olumayowa/Temporal_Domain_Multiplexing.git

https://gitlab.wllsb.edu.cn/blank22/kiryl-piatkevich/-/tree/master/Alex/Temporal%20domain%20multiplexing/Results

### Competing Interest Statement

The authors have declared no competing interest.

